# Label-Free Hyperspectral Imaging and Deep-Learning Prediction of Retinal Amyloid β-Protein and Phosphorylated Tau

**DOI:** 10.1101/2022.06.03.494650

**Authors:** Xiaoxi Du, Yosef Koronyo, Nazanin Mirzaei, Chengshuai Yang, Dieu-Trang Fuchs, Keith L. Black, Maya Koronyo-Hamaoui, Liang Gao

## Abstract

Alzheimer’s disease (AD) is a major risk for the aging population. The pathological hallmarks of AD—an abnormal deposition of amyloid β-protein (Aβ) and phosphorylated tau (pTau)—have been demonstrated in the retinas of AD patients, including in prodromal patients with mild cognitive impairment (MCI). Aβ pathology, especially the accumulation of the amyloidogenic 42-residue long alloform (Aβ_42_), is considered an early and specific sign of AD, and together with tauopathy, confirms AD diagnosis. To visualize retinal Aβ and pTau, state-of-the-art methods use fluorescence. However, administering contrast agents complicates the imaging procedure. To address this problem, we developed a label-free hyperspectral imaging method to detect the spectral signatures of Aβ_42_ and pS396-Tau and predicted their abundance in retinal cross sections. For the first time, we reported the spectral signature of pTau and demonstrated an accurate prediction of Aβ and pTau distribution powered by deep learning. We expect our finding will lay the groundwork for label-free detection of AD at its very earliest roots.

**Significance Statement:** The pathological hallmarks of Alzheimer’s disease (AD), amyloid β-protein (Aβ) and hyperphosphorylated (p)Tau protein have been characterized by a hyperspectral camera in terms of spectral signatures. The unique optical properties of the hallmark proteins on the broad visible light range enable label-free and high-resolution detection and virtual staining of abnormal deposition in the retina tissue, which will lay the groundwork for AD early diagnosis and AD development quantification.

## Introduction

Alzheimer’s disease (AD) and associated dementia are estimated to afflict 50 million people worldwide, a number projected to triple by 2050. This age-dependent epidemic is a major concern for the aging population, with an incidence that rises sharply after 65 years of age, affecting roughly 50% of individuals aged 85 and older.^[1]^ While currently there is no cure, with early diagnosis, the progression of the disease may be slowed and the patient life style may be changed.^[2,3]^

Although AD has been historically perceived as a brain disorder, recent studies indicate that AD also manifests in the eye with mounting evidence of abnormalities in the retina, a sensory extension of the brain.^[4–6]^ Particularly, the hallmark pathological signs of AD, amyloid β-protein (Aβ) and neurofibrillary tangles (NFTs) comprised of hyperphosphorylated (p)Tau protein, which have long been described in the brain, have also been identified in the retina.^[5,7]^ There is a growing number of reports that Aβ deposits and pTau were discovered in the retinas of AD patients at various stages, in stark contrast to non-AD controls.^[5,6,8–16]^ As the only central nervous system (CNS) tissue not shielded by bone, the retina offers unique access to study pathological changes in the brain, noninvasively and with unprecedented high spatial resolution. The evidence of Aβ accumulation in the retina at early stages of AD ^[5,13]^ and the accumulation of retinal NFT and pTau^[6,8,12]^ lends credence to the notion of the eye as a site for presymptomatic stage imaging. Notably, Koronyo-Hamaoui group and other teams revealed that retinal Aβ plaques, Aβ oligomers and pTau tangeles in transgenic AD-model mice appear at the presymptomatic and early stage and prior to detection in the brain.^[5,17–19]^

Despite holding great promise for early diagnosis of AD, visualization of retinal Aβ and pTau deposits is non-trivial. Because Aβ and pTau deposits have a similar visual appearance to normal tissue, conventional fundus photography provides little contrast. To increase visibility, state-of-the-art methods use exogenous fluorophores, and they have visualized retinal Aβ and pTau deposits with a high resolution.^[5–7,15,20–28]^ However, administrating contrast agents in humans complicates the imaging procedure, hindering its scalability for population screening. To date, only curcumin, a natural fluorochrome, has been tested and used in clinical trials to label retinal Aβ,^[6,21–23,25]^ whereas fluorophores used to visualize retinal pTau in vivo are more limited. Therefore, there is an unmet need to develop label-free, high-resolution imaging techniques to visualize retinal Aβ and pTau deposits for early AD screening and disease management.

Over the past decade, hyperspectral imaging (HSI) has been increasingly used in various medical applications, and it has shown promising results for detecting various cancers,^[29–37]^ diagnosis of cardiac^[38,39]^ and retinal diseases,^[40–42]^ and assessment of brain functions and activities.^[43–45]^ The overall rationale of using HSI for medical imaging is that the tissue’s endogenous optical properties, such as absorption and scattering, change during the progression of the disease, and the spectrum of light emitted from tissue carries quantitative diagnostic information about tissue pathology. Rather than measuring only light intensities at a two-dimensional grid, HSI captures a series of images at different wavelengths and forms a three-dimensional datacube (*x,y,λ*) (*x,y*, spatial coordinates; *λ*, wavelength), also known as a hypercube. The rich spatio-spectral information obtained enables the classification of chemical constituents of the tissue without fluorescence labeling.

By virtue of its label-free imaging ability, several pioneer groups have explored HSI in examining the optical characteristics of Aβ in paired brain and retina tissues from both transgenic AD mouse models and human AD patients.^[16,46–52]^ It has been found that the effect of Aβ can be dictated by a characteristic light reflectance spectrum, and the magnitude of the spectrum varies with the AD development, in stark contrast to non-AD population where no evident differences are detected. However, the HSI experiments performed so far lack validation against fluorescence-staining ground truth images, and their methods are inadequate to reveal the precise locations and types of Aβ deposits on the retina. Moreover, despite being equally important in AD pathology, to our knowledge, the spectral signature of pTau and its label-free detection by HSI have not been reported.

In this paper, we present a quantitative study on HSI of Aβ and pTau deposits in human retinal cross sections from neuropathologically confirmed AD patients. For the first time, we identified the spectral signature of pTau and demonstrated an accurate prediction of amyloidogenic 42-residue long (Aβ_42_) alloform and pS396-Tau deposits in the retina by utilizing a deep-learning approach. The Aβ_42_ and pS396-Tau markers were selected due to their recognized role in AD pathogenesis. For validation, we compared HSI prediction results with peroxidase-based immunostaining (also referred to as DAB staining) and immunofluorescent staining on the same imaging sections, which are both gold standards in quantifying Aβ and pTau deposits in retinal tissues.^[6,8,12,13]^ By feeding the spatio-spectral features associated with Aβ_42_ and pS396-Tau into a generative adversarial network (GAN), our method can also transform a label-free HSI image to either a DAB or an immunofluorescent stained image with high fidelity. The work presented here, therefore, lays the foundation for using HSI for non-invasive early AD diagnosis.

## Results

### Detection of Retinal Aβ_42_ and pS396-Tau Spectral Signatures by HSI

Using a custom HSI microscope equipped with a liquid crystal tunable filter (detailed configuration in Methods), we imaged unstained postmortem retinal cross sections from neuropathologically confirmed AD patients in the transmission mode. The retina cross-sections were prepared undergone tissue isolation, processing, and sectioning of superior-temporal (ST) and inferior-temporal (IT) strips. The hypercubes obtained contain the spatio-spectral information of endogenous chromophores in the retinal tissues. To guide spectral profiling, we immunostained retinal cross-sections specifically against Aβ_42_ and pS396-Tau and labeled either with peroxidase-based DAB substrate (3,3’-diaminobenzidine; brown) or immunofluorescence and reimaged it under a brightfield or fluorescence microscope, respectively (Zeiss Axio Imager Z1). We further registered the unstained hyperspectral images with the immunolabelled DAB or fluorescently stained images (Methods) and located the enriched areas of Aβ_42_ and pS396-Tau in the hyperspectral images. The spectral signatures of retinal Aβ_42_ and pS396-Tau were identified by averaging the pixel spectra in those regions (Figure 1 and 2), where Aβ_42_ and pS396-Tau exhibit distinct spectral profiles (more results are available in Figure S2-S4). Noteworthily, although the spectrum of Aβ_42_ has been previously reported, this is the first time the spectrum of pS396-Tau is identified.

**Figure 1.**
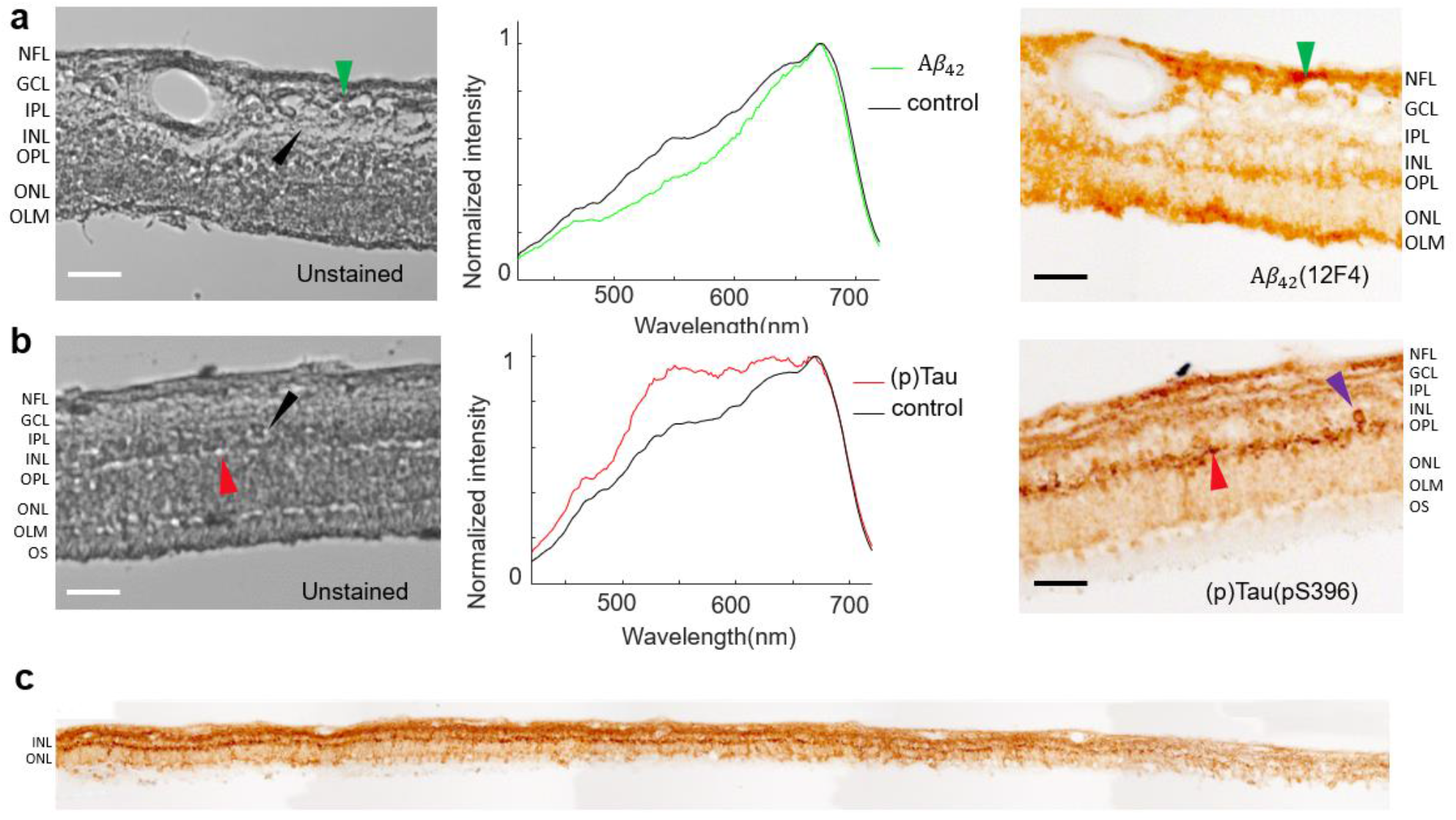
Hyperspectral imaging of **a**. 12F4^+^-Aβ_42_ and **b**. pS396-Tau deposits on postmortem retinal cross sections of AD patients. The Braak stages for patients are both V. AD patient in a is a female aged 90 with an ADNC score of A2, B3, C3. AD patient in b is a female aged 85 with an ADNC score of A3, B3, C3.From left to right, unstained hyperspectral intensity images, spectra at arrow-pointed locations (green for Aβ_42_, red for pS396-Tau, and black for control), and DAB labeled images. The purple arrow is indicating a NFT-like or cellular structure (b, right) in the OPL. **c:** A tile image of a large portion of retinal cross-section strip from a confirmed AD patient immunolabeled for pS396-Tau and DAB substrate. Scale bar of a and b, 50 µm.

**Figure 2.**
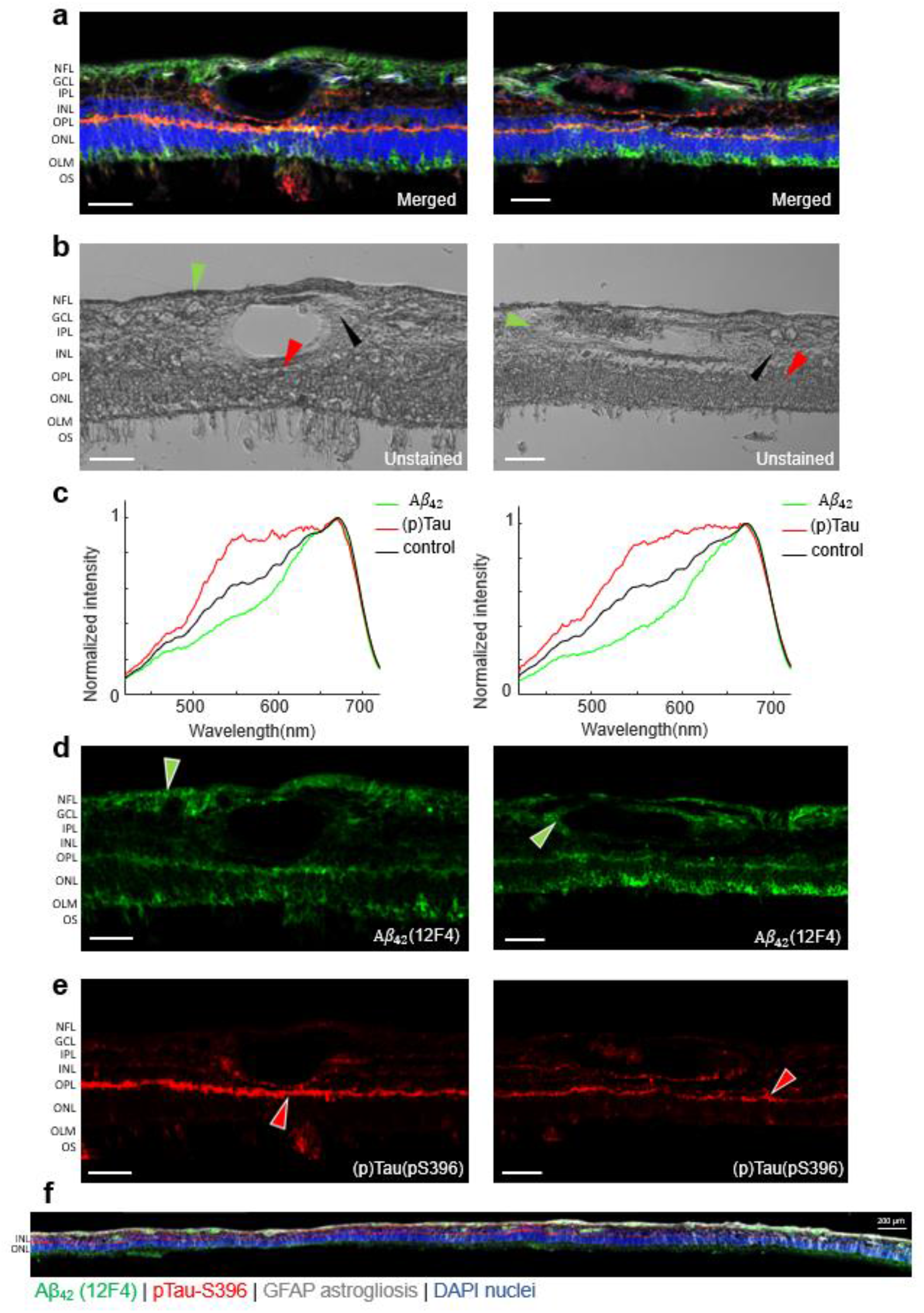
Hyperspectral imaging of Aβ_42_ and pS396-Tau deposits on postmortem retinal cross sections of AD patients guided by immunofluorescence staining. **a**. Merged fluorescence images of four channels. **b**. Unstained hyperspectral intensity images. **c**. Spectral signatures of Aβ_42_ and pS396-Tau in the human retina confirmed by combined fluorescence staining specific for 12F4^+^-Aβ_42_ and pS396-Tau. **d**. Aβ_42_ channel of a with green pseudo color. **e**. pS396-Tau channel of a with red pseudo color. **f**. A tile image of a large portion of retinal cross-section strip from a confirmed AD patient (Female, Age: 90, Braak stage: V, ADNC: A2, B3, C3) immunolabeled with combination antibodies against Aβ_42_ (green), pS396-Tau (red), and GFAP-astrocytes (white), and nuclei counterstained with DAPI (blue). Scale bar, 50 µm.

Consistent with previous studies,^[6,7]^ the immunolabeling with DAB or immunofluorescence-stained images indicate that pTau mostly aggregates in the retinal outer plexiform layer (OPL), inner plexiform layer (IPL), and ganglion cell layer (GCL), and in structures that resemble neurofibrillary tangles (NFTs) (Figure 1b,c). We also found pS396-Tau in the innermost retinal layers, along the nerve fiber layer (NFL), though it is variable from patient to patient and generally to a lesser extent (Figure S3a). We examined these locations in the unstained HSI images. Figure 1c shows the distribution of pS396-Tau deposit from the central to peripheral retina.The pS396-Tau clusters exhibit a unique spectral profile that significantly differs from that of ‘normal’ retinal tissues—they have a much higher and uniform transmittance for light in the 550 ∼650 nm range, resembling a “flat hat”. This dominating feature indicates that pTau enriched tissues have a reduced optical density in this spectral range, likely due to a smaller absorption coefficient of constituent chromophores. This prompted us to further examine the HSI images at these wavelengths. We found that the pTau aggregated in the OPL—which appears dark brown with DAB substrate and red in the immunofluorescence-stained images—correlates with higher pixel intensities in the grey-level HSI images (Figure 1b and 2). A similar correspondence has also been identified in the pS396-Tau aggregated region in the NFL (Figure S3a), corroborating our finding on the spectral transmission property of pTau. Notably, this is the first demonstration of pS396-Tau in human retina.

Besides the spectrum of pTau reported above, we also observed the known spectrum of Aβ. The DAB-or immunofluorescent-stained images show that specifically 12F4^+^-Aβ_42_ is most abundant in the retinal NFL, GCL, OPL, and the outer nuclear layer (ONL; Figure 1a and 2a,d). Moreover, Figure 2f indicates that the important vascular Aβ_42_ distribution is typical along the retinal cross-section strip. The spectra extracted at these locations in HSI images (Figure 1a and 2b) show a lower transmission in the 450 – 600 nm range, which were hypothesized to be caused by an elevated level of Rayleigh scattering in Aβ_42_ enriched tissues.^[53]^ We validated the consistency of the spectrum of Aβ_42_ in across all retinal layers including within blood vessel walls (extended data in Figure S2 and Figure S4). Noteworthily, although the spectral signature of retinal Aβ has been previously reported, this is the first time retinal Aβ_42_ has been reported and quantified at locations verified by fluorescence and non-fluorescence ground truth.

### GAN Network for Retinal Histopathology Image Prediction

Using the spatio-spectral information in hypercubes, we can classify the HSI images and generate abundance maps of constituent components. The images so obtained can be further rendered to resemble DAB and immunofluorescence staining using a pseudo-colormap. Among the state-of-the-art HSI classification approaches, deep learning (DL) is the most attractive option because it is robust against noise.^[54,55]^ Conventional DL methods classify HSI images pixel-wise solely based on the pixel’s spectral information.^[56,57]^ However, this usually leads to unsatisfactory results due to the missing link to the spatial features. Later endeavors improve the model by classifying the images in patches, followed by mosaicking the resultant abundance maps.^[58,59]^ Nonetheless, the resultant classified images suffer from a low resolution, and it is challenging to form a histopathology-like image.

To solve these problems, we adapted a generative adversarial network (GAN) for HSI classification of Aβ and pTau and image transformation. GAN is a competitive network consisting of a generator and a discriminator. The discriminator network is trained to classify the real inputs and the fake inputs generated by the generator network. This adversarial training increases the generalization capability of the discriminator, and it is particularly effective when the training dataset is limited.

For HSI classification, it is important to combine the complex spectral information of every pixel with the neighboring pixels’ information in a considerably efficient way. Most spatio-spectral-based classification methods use only a small neighboring region to construct a spatio-spectral vector.^[60–62]^ Although this can improve the classification accuracy than extracting only spectral information, the classification accuracy is limited by the size of the selected region. In contrast, the GAN network in our method considers the spatio-spectral features in the entire region imaged to assign a value for each pixel.

We developed a workflow to transform unstained human retinal cross-sections into two types of standard histopathology images (immunofluorescence and DAB) (Figure 3). We first convert an acquired hypercube (*x,y,λ*) to a three-channel image by principal component analysis (PCA) to represent the significant differences of the imaged pixel spectra. This significantly reduces the data load for training while preserving most of the variability (>85%) in the original hypercube. We then pair this extracted three-channel HSI image patch with the corresponding histopathology image and pass this image pair to the adapted GAN network for training. After training, we have four models for Aβ_42_ and pS396-Tau with two staining contrasts. The transformed histopathology-like images are output from the models and stitched to form a meaningful ROI.

**Figure 3.**
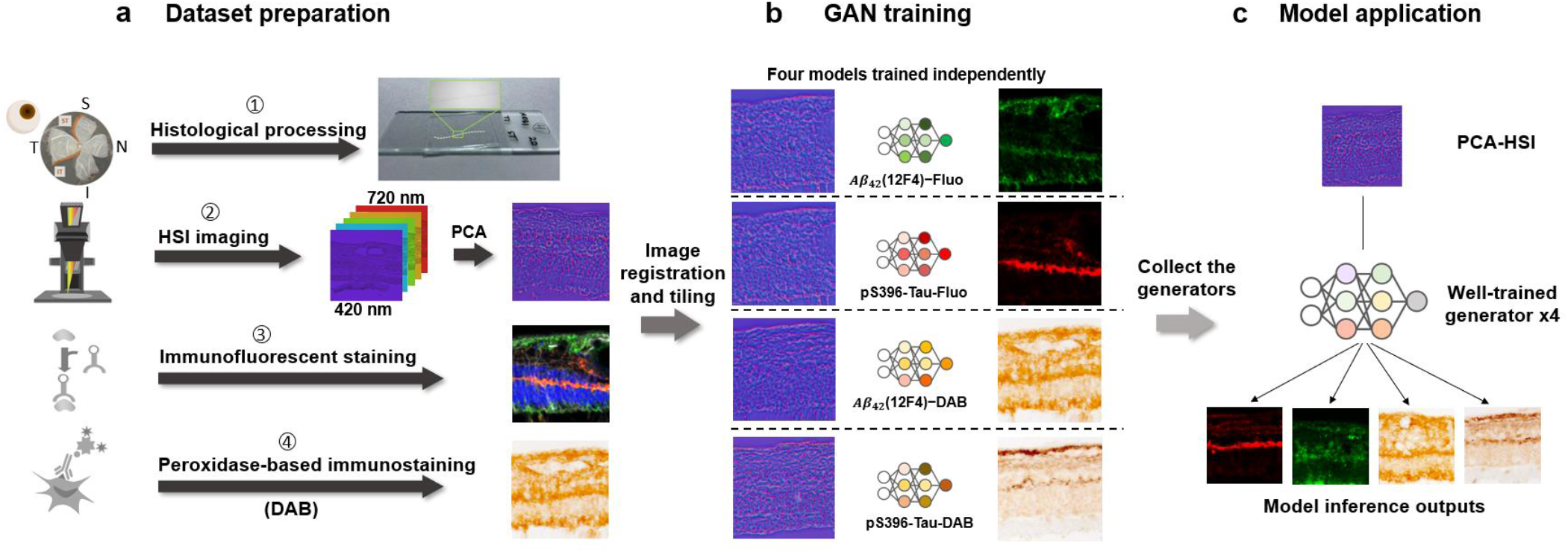
The deep learning workflow of Aβ_42_ and pS396-Tau deposits prediction. **a**. 1. The process of retina cross-sections preparation: donor eye fixation, neurosensory retina isolation to flatmounts, creating four retinal quadrants (S: superior, T: temporal, I: inferior, N: nasal), and sectioning of superior-temporal (ST) and inferior-temporal (IT) strips. 2. The retina cross-sections are imaged by our HSI microscope before immunostaining. Raw data are normalized and go through PCA process to convert to RGB images. 3 and 4 are two different staining techniques used for comparison of the HSI analysis. **b**. HSI images are registered with fluorescence and DAB staining images. Patches of size 256×256 pixels are cropped from the HSI and staining images. Corresponding images form training pairs for the generative adversarial network. **c**. We input the ROI patches to the trained model and get the inference patches for Aβ_42_ and pS396-Tau with different immune contrasts.

In immunofluorescent image transformation, we employed the green and red channels of the merged fluorescent images in Figure 2a as the ground truth for Aβ_42_ and pS396-Tau classification. In Aβ_42_ and pS396-Tau immunofluorescence stained (Cy5, green and Cy3, red pseudocolors, respectively) images, there were distinct signals for each marker and no autofluorescence signal in the lumen of blood vessels (Figure 2a, d-e). Although, autofluorescence signals were occasionally found at the blood vessel lumen (Figure 2f). Such signals will mislead the network training and prevent the true Aβ_42_ and pS396-Tau signals from forming proper contrast in the analyzed region. To solve this issue, we removed the lumen signals by labeling them as negative and enhanced the contrast of true Aβ_42_ and pS396-Tau signals (Figure S4). The models were trained by feeding green and red fluorescence data separately. The transformed immunofluorescent image patches of Aβ_42_ and pS396-Tau were stitched to a larger field-of-view (FOV) and shown in Figure 4a and 4b, respectively. In the zoomed insets, the predicted distributions of two AD-hallmark proteins in the region of interest match well with the ground truth. The Figure S6 shows the vascular wall Aβ_42_ deposit prediction with a negatively labeled lumen, and the Figure S7 shows various pS396-Tau deposits. In general, using immunofluorescence stained image as the ground truth yields an accurate prediction for Aβ_42_ and pS396-Tau deposit distribution. The specific signals in the actual histopathology image can also be identified in the transformed image. For instance, we found correspondence in the ground truth image for both the recovered Aβ_42_ signal pattern in Figure 4a and the transformed pS396-Tau in OPL image in Figure 4b.

**Figure 4.**
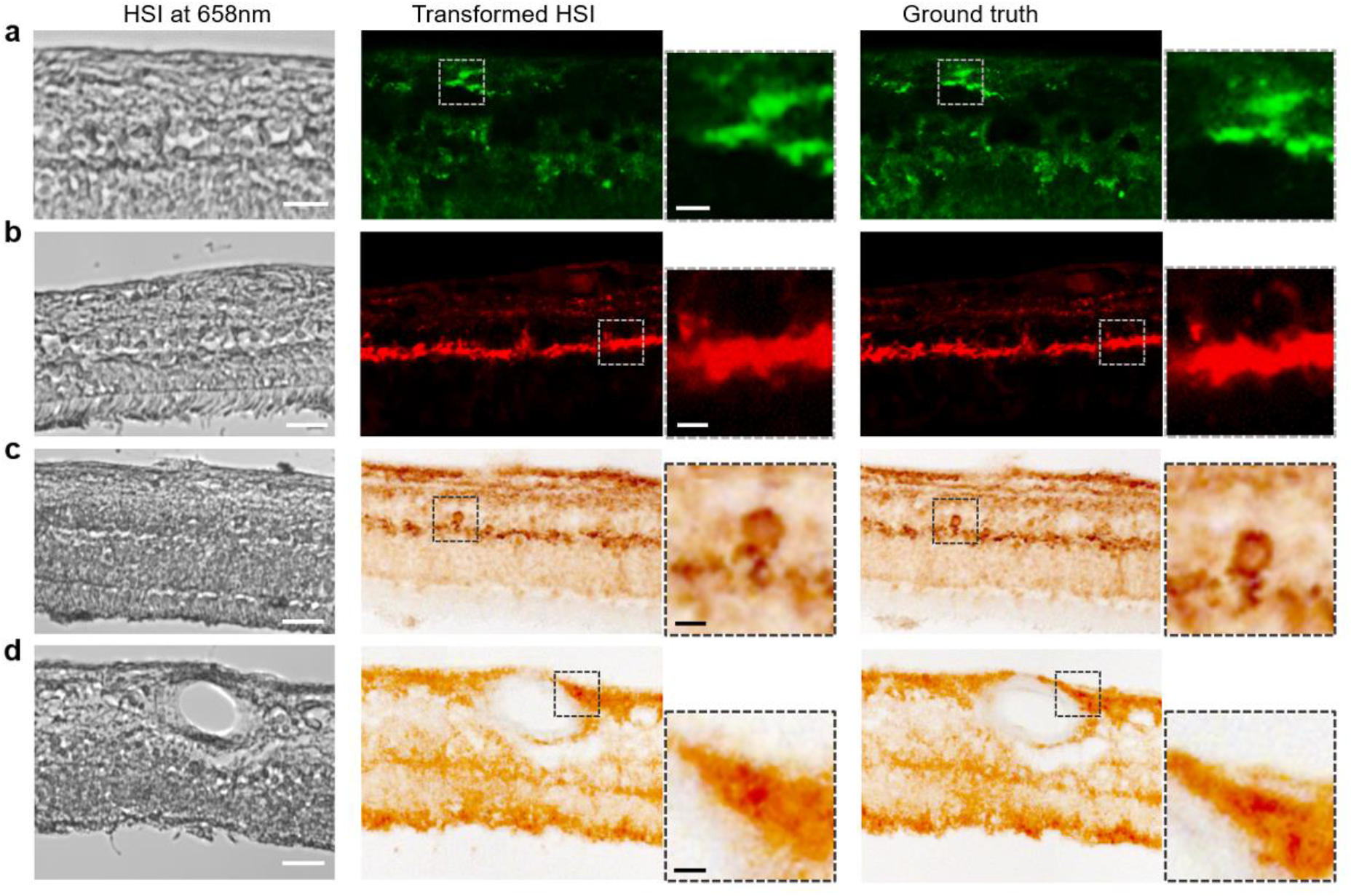
Stitched ROIs of the trained GAN models output. **a**. Aβ_42_ fluorescence model. **b**. pS396-Tau fluorescence model. **c**. pS396-Tau DAB model, with a focus on a retinal NFT structure. **d**. Aβ_42_ DAB model. From left to right: HSI intensity image, transformed HSI images, zoomed prediction images of specific feature, ground truth images, and zoomed ground truth images of specific feature. Scale bar, 50 µm for large FOV images, 10 µm for bordered inserts.

Besides immunofluorescence staining, we also stained the retinal cross-sections with the same primary monoclonal antibodies and using a highly sensitive immunoperoxidase-based DAB substrate.^[6,13]^ The DAB-stained retinas have only one channel for the specific labeled protein, and we imaged them using a bright field microscope. The image so obtained has accurate single protein contrast and provides a better view of tissue structures. For DAB image transformation, the networks were trained to assign classification values to pixels and learn the color scheme that appeared in DAB images. The trained DAB models can map the extent of Aβ and pTau deposits in a broad range with the DAB brown-color scheme. In Figure 4c, the transformed DAB-pS396-Tau image clearly shows layers of pTau deposits from the innermost layers to the OPL. In the OPL region, the structure of NFTs can be identified and visualized by our model (zoomed insert). There is also a signature band in the transformed DAB-pTau image, highlighting the pS396-Tau aggregation in the OPL with apparent deposit patterns. In some regions or patients, the inner retina has comparable pS396-Tau aggregates to that in the OPL, appearing in dark brown spots and most connected, as shown in Figure 4c and Figure S8. More results showing pS396-Tau OPL aggregation across retinal layers are provided in Figure S5c and S8. On the other hand, the DAB-Aβ (12F4 mAbs clone) images show that Aβ_42_ deposits appear in most retinal cell layers of cross-sections, a distribution that differs from pTau. The zoomed insert image in Figure 4d shows the predicted NFL/perivascular Aβ_42_ accumulation, a location where prominent Aβ_42_ signals have been found in our previous studies.^[6,13]^ Additionally, Aβ_42_ distributes in NFL and GCL to a large extent, which can be seen in other FOVs as well (Figure S9b-c, right). In this human cohort, Aβ_42_ deposits have also been found in the outer retina, especially in the ONL close to the outer limiting membrane, including the photoreceptor layer. Overall, Aβ_42_ in confirmed AD dementia patients usually presents in both the inner and outer layers of retinal cross-sections.

The GAN network is essential for learning the complex spectrum difference and distinguishing biomarkers in the microenvironment of retinal tissues, assessing their distributions, and potentially generating histopathology-like images to facilitate AD diagnosis. To build the model, we excluded the regions to be analyzed when preparing the training datasets. Also, to avoid inaccurate classification caused by the overlay of channels in immunofluorescence staining, we trained the network with the Aβ_42_ and pS396-Tau channels separately. We enlarged all datasets by data augmentation, mimicking the registration error between HSI and ground truth images (Methods), which, in turn, made the network more robust. The output prediction, a transformed histopathology-like image, correlates well with the ground truth. Compared with other spectral-based algorithms that take all spectral bands into training, our method requires less extensive computation, reduces network complexity, and makes training more efficient.

### Evaluation of the GAN Transformed Histopathology Images

We adopted a structural similarity index measure (SSIM) to assess the similarity between the transformed histopathology images by the GAN network and the actual stained images. SSIM is a perception-based image quality metric,^[63]^ which has been widely used to evaluate the structural similarities between synthesized images in deep-learning-based methods. SSIM equals one means a perfect match, whereas close to zero indicates hardly similar images.

Figure 5a shows the averaged SSIM values for the four trained models: DAB-pTau, DAB-Aβ, immunofluorescence-pTau, and immunofluorescence-Aβ. DAB-pTau has the highest SSIM value of 0.8714 (+/-0.0122), while immunofluorescence-Aβ has the lowest SSIM value of 0.8128 (+/-0.0219). Overall, the DAB models have a higher transformation performance than immunofluorescence models in our study when predicting the same biomarker deposits, and Aβ_42_ histopathology images have lower transformation accuracies than pS396-Tau in both stains. This is possibly due to the fact that Aβ_42_ is more abundant in the retina of AD patients than pS396-Tau, and it is highly dependent on disease development and assembly types, while pS396-Tau is found to be more layer-specific. Due to sample deformation during immunofluorescence staining, the HSI images could not be precisely registered to the immunofluorescence images, a fact that also lowers the SSIM values of immunofluorescence models.

**Figure 5.**
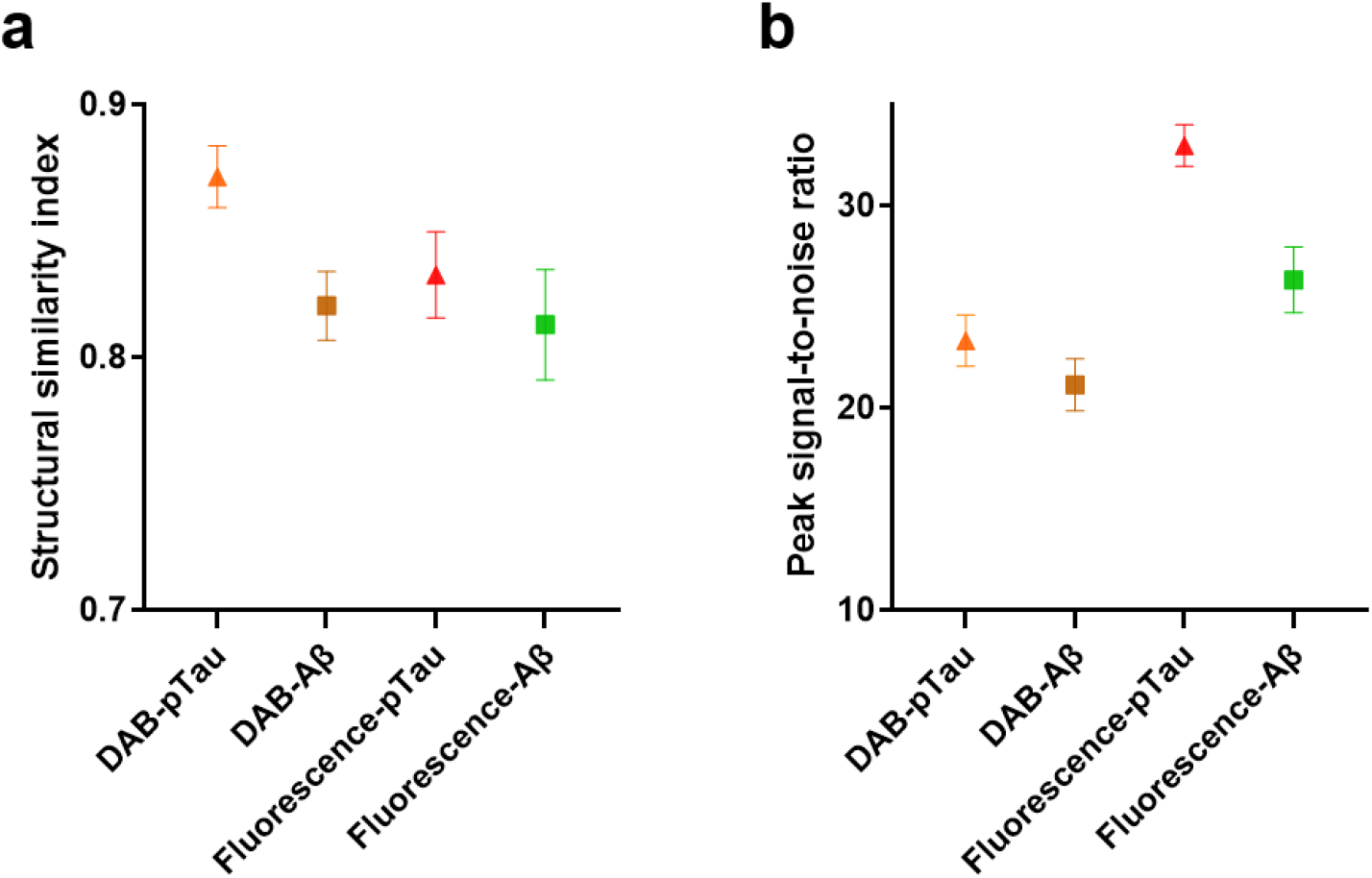
Averaged structural similarity index and peak signal-to-noise ratio (in dB) of the four models with error bars (standard deviation). DAB-pTau/Aβ: models used to transform HSI image to Peroxidase-based immunostaining image. Fluo-pTau/Aβ: models used to transform HSI image to immunofluorescence staining image.

The quantitative evaluation implies that the deep learning framework can transform the HSI images to histopathology images in high accuracy with a minimum SSIM of 0.8128. For the DAB-pTau model, we obtained an SSIM of 0.8714, which indicates the network can successfully recover the immunostaining color scheme and discriminate retinal pS396-Tau deposits. For comparison, a previous study that used a GAN network to transform quantitative phase images to H&E images achieved an SSIM value of only 0.80.^[64]^

In addition to SSIM, the peak signal-to-noise ratio (PSNR) was used as the second metric to evaluate the image quality of the transformed histopathology images (Figure 5b). The images generated by the immunofluorescence-pTau model have the highest PSNR of 32.9626 (+/-1.03) dB. The immunofluorescence-Aβ images have the second-highest PSNR of 26.3196 (+/-1.63) dB. The DAB-pTau and DAB-Aβ models have a PSNR value of 23.3136 (+/-1.27) dB and 21.1323 (+/-1.29) dB, respectively. All four models provide a PSNR value greater than 20 dB, indicating a high image quality. The immunofluorescence models have better image transformation quality than DAB models regarding PSNR. This might be because immunofluorescence stained images have a less complex color assignment and a black background compared with DAB staining. Notably, the two pTau transformation models still outperform Aβ models like that in the SSIM metric. A table summarizing all SSIM and PSNR values is provided in the supplementary information.

## Discussion

We demonstrated a hyperspectral imaging platform that enables label-free, high-resolution structural and molecular imaging of Aβ and pTau deposits in human retinas. The advantage of HSI is its label-free imaging ability by capturing tissue spectrum across a broad spectral range. Our system illuminates the sample using a simple broadband halogen lamp and scans the sample wavelength using a liquid crystal filter. The quantified intensity values from the spectral channels imply the optical characteristics of the AD biomarkers, Aβ and pTau. The discovered Aβ_42_ and pS396-Tau spectral signatures are highly consistent in various regions of retinal cross-sections and among the patients. We also examined the consistence of the HSI data by reimaging the samples in an extended period of time (14 days), and we found that the spectral signatures of both Aβ_42_ and pS396-Tau remain the same. More importantly, we are the first to report the pS396-Tau in the human retina and its spectral signature. We further visualize pS396-Tau with a label-free imaging technique. Our method, which can probe pTau deposits, has the potential to advance AD quantification and diagnosis. Seeing its label-free imaging ability and system simplicity, we also anticipate the presented HSI method will become an alternative or complementary approach to histopathological analysis of Aβ and pTau in CNS organs and other tissues.

Our findings on the spectral signature of Aβ echo many pioneer works in the field. For example, Xavier Hadoux et al.^[16]^ discovered a significant difference in ocular reflectance among patients with and without moderate-to-high Aβ levels, and they confirmed their findings through imaging the paired brain samples. As another example, More’s group^[46–48]^ reported the spectral signature of Aβ_42_ in both retina and brain tissues in human and transgenic mouse. Nonetheless, all these previous studies lack a direct validation through immunostaining. Moreover, because the measurements were performed in the widefield imaging mode, the distribution of Aβ in retinal cross sections remains elusive. Our findings presented herein, therefore, provides the basis for the previous research.

In addition, we developed a framework to facilitate the classification of the spectral signatures of retinal Aβ_42_ and pS396-Tau and transformed the unstained HSI images to histopathology-like images.

Our GAN network is robust to local misalignments in registration and staining overflow because we created augmented training data to mimic those influences. This is especially useful for immunofluorescent image transformations because the staining process is more complex and less specific than the DAB-staining. The inaccuracies in the staining and image acquisition processes must be taken into consideration to avoid confusing the network. Another benefit of using the transformation framework is that it can be trained to generalize the variations of the histopathology stained tissues across different sections and patients with sufficient datasets.

The Aβ_42_ and pS396-Tau in the two standard staining techniques, DAB and immunofluorescence, were trained separately as four models. Multi-channel immunofluorescence transformed images of a single tissue region can be achieved by combining the generated Aβ_42_ and pS396-Tau images in green and red channels. An advantage of training separately, especially for immunofluorescence stains, is avoiding overlapping between two channels. With a well-trained transformation model, the histopathology images can be generated instantaneously, without the need for tedious pathological processing.

In conclusion, we developed a label-free HSI method as a tool to report Aβ_42_ and pS396-Tau spectral signatures and a deep-learning-based framework to transform the unstained HSI image to a histopathology-like image. Our method thus democratizes the immunofluorescence/DAB staining and makes them accessible to general labs. Also, our entire workflow (Figure 3) is time-efficient. Scanning the sample and computing the transformed images take less than 30 min, which is only a fraction of the time typically needed when the sample undergoes the conventional pathological processing (2-4 days).

We expect our method will lay the foundation for future label-free AD screening and diagnosis using HSI approaches.

## Methods

### Human Eye Donors

Postmortem human eyes were obtained from the Alzheimer’s Disease Research Center (ADRC) Neuropathology Core in the Department of Pathology (IRB protocol HS-042071) of Keck School of Medicine at the University of Southern California (USC, Los Angeles, CA). USC-ADRC maintains human tissue collection protocols that are approved by their managerial committees and subject to oversight by the National Institutes of Health. Histological studies at Cedars-Sinai Medical Center were performed under IRB protocols Pro00053412 and Pro00019393. For the histological examination, 12 retinas were collected from deceased patient donors. The retinas from 10 donors with clinically and neuropathologically confirmed AD (n=2), MCI (n=3), and cognitively normal (CN; n=5), were used in the early stage of the training phase of immunostaining (demographic data on human donors are given in Table S3). Addionally, n=3 neuropathologically confirmed AD dementia patients were used for histological and HSI analyses followed by network training; donors’ age, gender, ethnic background, premortem and final diagnosis, Braak stage, CDR and/or MMSE score and postmortem interval (PMI) of tissue collection are detailed in Table S4. All samples were de-identified and could not be traced back to tissue donors.

### Clinical and Neuropathological Assessments

The USC ADRC Clinical Core provided clinical and neuropathological reports on the patients’ neurological examinations, neuropsychological and cognitive tests, family history, and medication lists. Most cognitive evaluations had been performed annually and, in most cases, less than 1 year prior to death. Cognitive testing scores from evaluations made closest to the patient’s death were used for this analysis. Two global indicators of cognitive status were used for clinical assessment: the Clinical Dementia Rating (CDR scores: 0 = normal; 0.5 = very mild dementia; 1 = mild dementia; 2 = moderate dementia; or 3 = severe dementia)^[65]^ and the Mini-Mental State Examination (MMSE scores: normal cognition = 24–30; MCI = 20–23; moderate dementia = 10–19; or severe dementia ≤ 9).^[66]^ In this study, the composition of the clinical diagnostic group (AD, MCI, or CN) was determined by source clinicians based on findings of a comprehensive battery of tests including neurological examinations, neuropsychological evaluations, and the aforementioned cognitive tests. To obtain a final diagnosis based on the neuropathological reports, we used the modified Consortium to Establish a Registry for Alzheimer’s Disease (CERAD)^[67]^ as outlined in the National Institute on Aging (NIA)/Regan protocols with revision by the NIA and Alzheimer’s Association.

### Preparation of Retinal Cross-Sections

Fresh-frozen eyes and eyes preserved in Optisol-GS were dissected with anterior chambers removed to create eyecups. Vitreous humor was thoroughly removed manually. Retinas were dissected out, detached from the choroid, and flatmounts were prepared.^[13]^ By identifying the macula, optic disc, and blood vessels, the geometrical regions of the four retinal quadrants were defined with regard to the left and the right eye. Flatmount strips (2–3 mm in width) were dissected along the retinal quadrant margins to create four strips: superior-temporal—ST, inferior-temporal—TI; inferior-nasal—IN, and superior-nasal—NS, and were fixed in 2.5% PFA for cross-sectioning. Each strip was approximately 2–2.5 cm long from the optic disc to the ora serrata and included the central, mid, and far retinal areas. All the above stages were performed in cold phosphate-buffered saline (PBS) with 1 × Protease Inhibitor cocktail set I (Calbiochem 539,131). Eyes that were initially fixed in 10% NBF or 2.5% PFA were dissected to create eyecups, and the retinas were dissected free. Vitreous humor was thoroughly removed and flatmounts were prepared. As described above, a set of flatmount strips, Superior Temporal (ST), Inferior Temporal (IT), Inferior Nasal (IN) and Nasal Superior (NS), was dissected (2–3 mm in width), washed in 1x PBS, and processed for retinal cross-sectioning.

Flatmount strips were initially embedded in paraffin using standard techniques, then rotated 90° horizontally and embedded in paraffin. The retinal strips were sectioned (7–10 µm thick) and placed on microscope slides that were treated with 3-Aminopropyltriethoxysilane (APES, Sigma A3648). Before immunohistochemistry, the sections were deparaffinized with 100% xylene twice (for 10 min each), rehydrated with decreasing concentrations of ethanol (100–70%), and then washed with distilled water followed by 1x PBS.

### Hyperspectral imaging

We built a hyperspectral imaging system based on an Olympus IX83 microscope. The samples are illuminated by a broadband halogen lamp, and the transmitted light is collected by a 10× objective lens (Olympus, 0.25 NA). The output image is filtered by a liquid crystal tunable filter (KURIOS-VB1, Thorlabs) in narrow bandwidth setting (10 nm FWHM at λ = 550 nm). The spectral range is from 420 nm to 720nm, with a wavelength scanning step of 2 nm. We collected the image data using a monochrome sCMOS camera (CS2100M, Thorlabs). In total, 151 spectral images were captured for one field-of-view (FOV). The entire cross section of the retina ST was scanned with a 1/3 overlap between adjacent FOVs for image stitching. A sample not in imaging was attached to a glass slide without a cover glass and kept in PBS 1x solution. When performing imaging, we placed a cover glass on top of the sample and replenished it with PBS 1x solution to keep the tissue moist. All retinal cross-section samples were kept in PBS 1x solution over two weeks and reimaged multiple times. Upon completion of scanning, we stitched all the FOVs at the selected wavelength to a whole strip view of the retina cross section.

### Light Source Calibration

Because the sCMOS camera has different spectral responses to different wavelengths, the acquired HSI data must compensate for the system response. We used a benchmark fiber spectrometer (O STS-VIS-L-25-400-SMA, Ocean Optics) to measure the lamp spectrum at the sample stage and imaged the slides with a blank FOV. The calibration coefficients for all the spectral components were obtained by dividing the average image intensities by measured spectral values. The calibration coefficients were fine-tuned by imaging a color checker (X-Rite Color Checker). The final calibration coefficients were saved and used in the following HSI data processing.

### HSI Data Processing

The spectral signature of retinal tissue was examined in an average manner by area, consisting of a minimum of three pixels. Each raw HSI tiff stack file contains 151 image slices. All the slices were read in and formed into a data cube format for efficient processing. The intensity at each spectral band was averaged over the selected area. Then the intensity values were calibrated by the pre-calculated calibration coefficients. The overall intensity of the imaging spectral range was normalized. We used the stained most adjacent slides (5μm distance) as a guidance and scanned the correspondence neighboring areas with marked deposits to reveal the spectral signature of Aβ_42_ and pTau. Control regions were selected as regions from the immunostaining images with neither Aβ_42_ nor pTau deposits and without cellular structures.We also scanned across the tissue vertically and horizontally to locate characteristic areas and plotted all the spectra. Then we classified the spectrum into several categories.

They were matched with the occurrence coordinates. The spectral graphs in Figure 1 and 2 are representative examples of the reported Aβ_42_ and pTau spectral signatures. More representative spectra graphs of various retinal locations can be found in supplementary information.

### Immunofluorescent Staining of Retinal Cross-Sections

After deparaffinization, retinal cross-sections were treated with antigen retrieval solution at 99 °C for 1 h (PH 6.1; Dako #S1699) and washed in 1x PBS. Retinal sections were then incubated in blocking buffer (Dako #X0909) and adding 0.2% Triton X-100 (Sigma, T8787), for 1h at RT, followed by primary antibody incubation overnight in 4 °C with the following combination: mouse anti-Aβ 1-42 antibody, 12F4 (1:500, BioLegend #805502) and rabbit anti-pTau antibody, pSer396 (1:2500, AS-54977). The 12F4 antibody is specific to the detection of amyloid beta x-42, without cross reacting with amyloid beta x-40 or amyloid beta x-43. Retinal sections were then washed three times by 1x PBS and incubated with secondary antibodies (1:200; Cy5 conjugated donkey anti mouse and Cy3 conjugated donkey anti rabbit, Jackson ImmunoResearch Laboratories) for 2 h at RT. After rinsing with 1x PBS three times, sections were mounted with Prolong Gold antifade reagent with DAPI (Thermo Fisher #P36935). Fluorescence images were repeatedly captured at the same focal planes with the same exposure time using a Carl Zeiss Axio Imager Z1 fluorescence microscope with ZEN 2.6 blue edition software (Carl Zeiss MicroImaging, Inc.) equipped with ApoTome, AxioCam MRm, and AxioCam HRc cameras. Tiling mode and post-acquisition stitching were used to capture and analyze large areas. Multi-channel image acquisition was used to create images with multiple channels. Images were captured at 20 ×, 40 ×, and 63 × objectives for different purposes. Routine controls were processed using identical protocols while omitting the primary antibody to assess nonspecific labeling.

### Peroxidase-Based Immunostaining of Retinal Cross-Sections

Retinal cross-sections after deparaffinization were treated with target retrieval solution at 99 °C for 1 h (pH 6.1; Dako #S1699) and washed with 1x PBS. In addition, treatment with 70% formic acid (ACROS) for 10 min at RT was performed on retinal cross-sections before staining for Aβ and pTau. For a list primary antibodies and dilutions, see above under Immunofluorescent staining. Following the treatment with formic acid, the tissues were washed with wash buffer 1x (Dako S3006) and adding 0.2% Triton X-100 (Sigma, T8787), for 1h, then were treated with 3% H2O2 for 10 min and washed with wash buffer. Each primary antibody was diluted with background reducing components (Dako S3022) and incubated separately with the tissues overnight in 4 °C. Tissues were rinsed with wash buffer three times for 10 min each on a shaker, then incubated separately for 30 min at 37 °C with secondary Ab (anti-mouse ab HRP conjugated, DAKO Envision K4001 or anti-rabbit ab HRP conjugated, K4003). Next, tissues were rinsed with wash buffer three times for 10 min each on a shaker. Liquid DAB+ Substrate Chromogen System (DAKO K3467) was used, then slides were immersed in dH2O and washed with wash buffer for 5 min, then washed with slow running tap water for another 5 min. Tissues were mounted with Faramount aqueous mounting medium (Dako, S3025). Routine controls were processed using an identical protocol while omitting the primary antibodies to assess nonspecific labeling. Brightfield images were repeatedly captured at the same focal planes with the same exposure time using Carl Zeiss Axio Imager Z1 microscope equipped with AxioCam HRc camera. Images were captured at 20 ×, 40 ×, and 63 × objectives for different purposes. Tiling mode and post-acquisition stitching were used to capture and analyze large areas.

### Image Registration

The retinal HIS cross-sectional images were co-registered with the immunofluorescence/DAB stained images of the same tissue section, for the purpose of analyzing the spectral signature and forming training pairs as the ground truth images for the transformation framework training. One spectral channel image with the most contrast was selected, and registered with immunostained images using affine transformation. The intensity values of immunofluorescence images (8-bit) were subtracting 255 from each image pixel to get a complement image. First, apparent tissue features such as blood vessels and edges are used to crop the corresponding immunostained images to the same FOV of HIS images. Then rotation, translation, and scaling operations are applied on immunostained images to produce a non-reflective similarity transformation. In cases that the affine registration is not sufficient by visual inspection, an optional control-point registration is applied using control point pairs selected from the tissue, such as blood vessel edges. The control point registration is implemented by a local weighted mean of inferred second-degree polynomials from each neighboring control point pair to create a transformation mapping.

### GAN Models

The GAN network used in our study is adapted from a conditional GAN.^[68]^ The generator network is based on a U-net using Pytorch. We incorporated a structural similarity index (SSIM) component into the generator loss function as −*v*× Log [(1 + *SSIM*(*G*(*x*),*y*))/2]. Mean absolute error (*L*_1_) loss is used to regularize the generator to transform the input image accurately and in high resolution. SSIM is used to balance the *L*_1_ loss of learning correct features rather than the pixel accuracies. The loss function has the following form:

Generator:

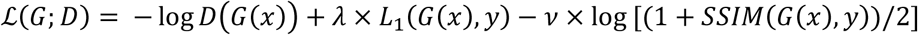

Discriminator:

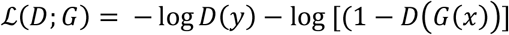

where *x* is the PCA-compressed HSI image, y is the ground truth image, and G/D denotes the forward pass of the generator/discriminator network, *λ* and *v*are weights to control the loss of *L*_1_ and SSIM terms. The network utilizes both spatial and spectral information to classify Aβ/pTau. The transformed histopathology images output from the generator were evaluated by a three-layer discriminator. We trained the compressed HSI images of Aβ and pTau and the two corresponding immunostainings separately and obtained four trained models. The four models were used to transform the test HSI images to Aβ and pTau stained immunofluorescence and DAB images. The weights for the loss function components were set as 100 for L1 loss, and 100 for SSIM term. We achieved optimal results with learning rates of 5× 10^−6^for the immunofluorescence models and 1 × 10^−5^for the DAB models using the adaptive moment estimation (Adam) optimizer. The batch size was set to one under the instance normalization. The epoch number was in between 120 and 150, with 50 epochs for decayed learning rate. Training time was approximately 47 h for immunofluorescence models and 82 h for DAB models. The network was implemented with one GTX TITAN graphical processing unit (GPU) using Pytorch 1.6.0, and Python version 3.6.8 on a desktop installed with Ubuntu 16.04 operating system. The desktop is equipped with CPU Intel Core i7-6900K@3.20 GHz and 64 GB RAM.

### Training data Preparation

The input format of our GAN network is an image pair consisting of one compressed HSI image and the corresponding ground truth image in 8-bit. We compressed the HSI data by applying PCA. The HSI image slices were cropped to only keep a small portion of the background next to the retinal tissue, to ensure most HSI data contained tissue spectral information. The first three principal components representing the spectral information were fed in order into the red, green and blue channels. The PCA compressed HSI image was cropped out as 256-pixel × 256-pixel × 3 patches, the corresponding ground-truth patches were cropped from the previous co-registered immunostained images. The selected analyzed regions of stained tissue were left and cropped as test image patches, with 1/3 to 1/2 overlap. This is to avoid the discontinuous intensities when stitching the transformed patches. The other parts of image were cropped and separated into training and validation set. Validation data was randomly selected from training data. The numbers of image pairs used for the four models training are given in Table S2. The patches containing damaged tissue regions and severely deformed staining tissue that leading to an unreliable registration during staining process were discarded.

### Data Augmentation

We implemented the traditional data augmentation techniques to enlarge our data size and make the network more robust to accommodate the offset that remained after the image registration. The operations include translation, rotation, flipping, scaling, and stretching, most of which were also applied in the registration process. These similar transformations make the network adapt to the registration offset. By generating data under those conditions, we increased our dataset size by a factor of 12 and improved the spatial criteria confidence.

### Image Stitching

The ground truth images, immunofluorescent staining and peroxidase-based immunostaining retinal cross-sections, were acquired at 20X using titling mode (multiple focus points were set) and stitched by the image stitching tool on Zen Blue Software to capture and analyze the entire retinal strip. For HSI retinal cross-section images, we chose one image slice at one well-contrasting wavelength and stitched all connected FOVs using the Image Composite Editor software. We stitched the output images from the trained GAN models with a self-derived algorithm. The algorithm iterates to find the best connective coordinate by scanning the corresponding overlap region of the two adjacent image patches, then stitches the images at this position. For the several cases when images have connective artifacts, we averaged the intensities of neighboring columns of the connective coordinate.

### Quantification Metric

The model output transformation images are compared to the corresponding ground truth using SSIM index and PSNR as similarity and quality measures. SSIM compares the transformed image with ground truth images in three measurements: luminance, contrast and structure. PSNR is a common tool to assess the image reconstruction quality and it is used to assess the compression ratio of the transformed image. For each output image patch, an SSIM value and a PSNR value were calculated. The average and standard deviation values were calculated for each model group. The SSIM metric is calculated between the transformed image *i*and the ground truth image *j*as:

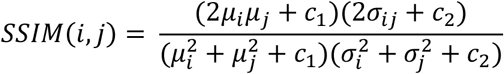

Where *μ*_*i*_and *μ*_*j*_are the averages of *i*and *j*; σ_*i*_and σ_*j*_are the standard deviations of *i*and *j*; σ_*ij*_is the covariance of *i*and *j*; and *c*_1_and *c*_2_ are regularization constants to avoid instability when the other variables are close to zero.

## Supporting Information

See for supporting contents.

## Acknowledgements

We thank the USC-ADRC Neuropathology Core and Dr. Carol Miller, MD, for providing access to human donor tissues. This article is dedicated to the memory of Dr. Salomon Moni Hamaoui and Lillian Jones Black, both of whom died from Alzheimer’s disease.

## Author Contributions

X.D. designed the system, built the system, performed the experiments, and analyzed the data. L.G., M.K-H. study conception and design. Y.K., D-T.F. collected and selected human eye globes, isolated the neurosensory retina and prepared retinal cross-sections. Y.K., N.M stained the retinal samples, and scanned and processed stained sample images. M.K-H., Y.K., N.M. identified and verified biological targets of interest in the AD retina. All authors contribute to the writing or editing of the manuscript. M.K-H., L.G. supervised the project.

## Code Availability

The codes used to perform the retina histopathological transformation in this manuscript can be found on Github at https://github.com/carinaxiaoxi/rcs_transform.

## Data Availability

The HSI data, immunofluorescent and DAB staining images supporting this study in the main text and supporting information are available on Github at https://github.com/carinaxiaoxi/rcs_transform.

